# Cell-to-cell spread inhibiting antibodies constitute a correlate of protection against Herpes Simplex Virus Type 1 reactivations

**DOI:** 10.1101/2020.09.03.282483

**Authors:** Susanne Wolf, Mira Alt, Robin Dittrich, Miriam Dirks, Leonie Schipper, Ulrich Wilhelm Aufderhorst, Kordula Rainer, Ulf Dittmer, Oliver Witzke, Bernd Giebel, Mirko Trilling, Christiane Silke Heilingloh, Ramin Lotfi, Michael Roggendorf, Adalbert Krawczyk

## Abstract

Herpes simplex viruses (HSV) cause ubiquitous human infections. For vaccine development, knowledge concerning correlates of protection against HSV is essential. Therefore, we investigated if humans principally can produce highly protective cell-to-cell spread inhibiting antibodies upon natural infection and whether such antibody responses correlate with protection from HSV reactivation. We established a high-throughput HSV-1 GFP reporter virus-based assay and screened 2496 human plasma samples for HSV-1 cell-to-cell spread inhibiting antibodies. We conducted a survey among the blood donors to analyze the correlation between the presence of cell-to-cell spread inhibiting antibodies in plasma and the frequency of HSV reactivations. In total, 128 of 2496 blood donors (5.1 %) exhibited high levels of HSV-1 cell-to-cell spread inhibiting antibodies in the plasma. Such individuals showed a significantly lower frequency of HSV reactivations compared to subjects without sufficient levels of HSV-1 cell-to-cell spread inhibiting antibodies. This study provides two important findings: (I) a fraction of humans produce HSV cell-to-cell spread inhibiting antibodies upon natural infection and (II) such antibodies correlate with protection against recurrent HSV. Moreover, these elite neutralizers can provide promising material for hyperimmunoglobulin, the isolation of superior antiviral antibodies and information for the design of a vaccine against HSV.

**Importance:** Herpes simplex virus 1 infections can cause painful mucosal lesions at the oral or genital tract and severe, life threatening disease in immunosuppressed patients or neonates. There is no approved vaccine available, and the emergence of drug resistances especially in long time treated patients makes the treatment increasingly difficult. We tested 2496 people for HSV-1 cell-to-cell spread inhibiting antibodies. Five percent exhibited functional titers such antibodies and showed significantly lower risk of reactivations, uncovering cell-to-cell spread inhibiting antibodies as a correlate of protection against Herpes simplex virus reactivations. Isolation of the cell-to-cell spread inhibiting antibodies from B-cells of these donors may contribute to develop novel antibody-based interventions for prophylactic and therapeutic use and provide starting material for vaccine development.

## Introduction

Herpes simplex viruses (HSV) types 1 and 2 are among the most common human infections worldwide. Globally, more than 3.7 billion people are infected with HSV-1 [1] and nearly 500-million with HSV-2 [2]. Both viruses cause a broad range of disease manifestations ranging from painful and irritating but self-limiting oral or genital lesions to severe disseminated and life-threatening infections in immunocompromised patients [2–5]. Serious complications can also be observed in patients suffering from ocular herpes infections, which may result in irreversible damage of the eye or even blindness [6, 7].

Until today, there is no approved vaccine available [8]. Numerous animal studies investigating the efficacy of distinct vaccine candidates such as inactivated virus particles, live- or genetically attenuated viruses or recombinant subunit vaccines yielded promising results [9, 10]. However, none of the vaccine candidates being tested in clinical trials has been effective [8]. The GlaxoSmithKline (GSK) Herpevac trial using a recombinant HSV-2 glycoprotein D (gD2) subunit vaccine was largest clinical trial performed so far [11]. In strong contrast to prior animal studies, the vaccine failed to protect against the acquisition of HSV-2 infection [11]. The discrepancy between promising results of animal studies and the failure of clinical trials in humans indicated a fundamental difference in the immune response to HSV in mice or guinea pigs and humans. Retrospective studies uncovered differences in antibody responses between humans and rodents concerning virus-specific antibodies, neutralizing antibodies, and cell-to-cell spread inhibiting, neutralizing antibodies (CCSi-NAbs). Most recently, the antibody responses to the gD2 subunit vaccine were analyzed in humans and guinea pigs [12]. Antibodies produced by vaccinated humans recognized significantly fewer crucial gD2 epitopes as compared to guinea pig antibodies [12, 13]. The crucial gD2 epitopes are targets of neutralizing or cell-to-cell spread inhibiting antibodies [14]. The cell-to-cell spread of HSV is known as a mechanism of immune evasion, and markedly facilitates the spread of HSV upon reactivation [14]. Antibodies, which can block this route of viral transmission, are associated with protection from disease [12, 15]. We developed a highly neutralizing and cell-to-cell spread inhibiting monoclonal antibody called mAb 2c. This antibody mediates almost complete protection from lethal genital HSV-1 infection - even in highly immunodeficient NOD/SCID mice [15, 16]. Moreover, mAb 2c protects mice from the development of severe ocular infections [17–19]. Importantly, mAb 2c is significantly more effective in protecting from disease than polyclonal human neutralizing antibodies used at a similar neutralizing titer, highlighting the importance of the inhibition of cell-to-cell spread in protecting from disease [20]. These *in vitro* and *in vivo* data strongly suggested that neutralizing antibodies, which inhibit the cell-to-cell spread are superior to antibodies that “just” neutralize but do not inhibit the cell-to-cell spread [20]. These findings raise the apparent question, if the inhibition of the cell-to-cell spread might contribute to protection from primary and/or recurrent disease. Intriguingly, the re-evaluation of the GSK Herpevac trial revealed that gD2 immunized individuals only barely produced antibodies that targeted gD2 epitopes associated with cell-to-cell spread [13], raising the fundamental question whether humans are in principle able to produce cell-to-cell spread inhibiting antibodies against HSV.

To address this question, we established a HSV-1 GFP reporter virus-based high-throughput screening assay and tested 2496 plasma samples for cell-to-cell spread inhibiting antibodies. We show for the first time that a small fraction of humans is indeed able to produce functional levels of cell-to-cell spread inhibiting antibodies (elite responder) and - even more striking - that the presence of sufficient concentrations of such antibodies correlated with protection from HSV reactivation.

## Results

To evaluate whether humans are able to produce potent antiviral antibodies upon natural HSV-1 infection, we established a high-throughput assay to test human plasma and serum samples for HSV-1 cell-to-cell spread inhibiting properties.

### HSV-1-ΔgE-GFP reporter virus-based screening assay for cell-to-cell spread inhibiting antibodies

The screening assay is based on the quantification of the progressing plaque expansion, which correlates with the extent of cell-to-cell spread. By using the HSV-1-ΔgE-GFP reporter virus, plaque formation, which is proportional to the GFP expression level, can be quantified using a fluorescence reader and visualized by fluorescence microscopy (Fig. 1). Confluent Vero cell monolayers were infected with the HSV-1-ΔgE-GFP reporter virus at a low multiplicity of infection (MOI = 0.001). Thereby, only scattered cells within the cell layer become infected. Afterwards, the infected cell cultures were overlaid with medium containing the sample to be tested, e.g. serum, plasma or purified antibodies (Fig. 1). The test was evaluated 3 days after infection in a quantitative manner by assessing the GFP signal or in a qualitative manner by fluorescence microscopy (Fig. 1). Single infected cells accompanied by a low GFP signal represent a complete inhibition of the cell-to-cell spread. Unrestricted plaque formation and strong GFP signals at levels similar to those of the HSV-1 seronegative control were scored as no inhibition of the cell-to-cell spread. Small plaques and moderate GFP signals indicated the presence of cell-to-cell spread inhibiting antibodies in the sample, even if there was no complete inhibition of the cell-to-cell spread (Fig. 1).

**Figure 1:**
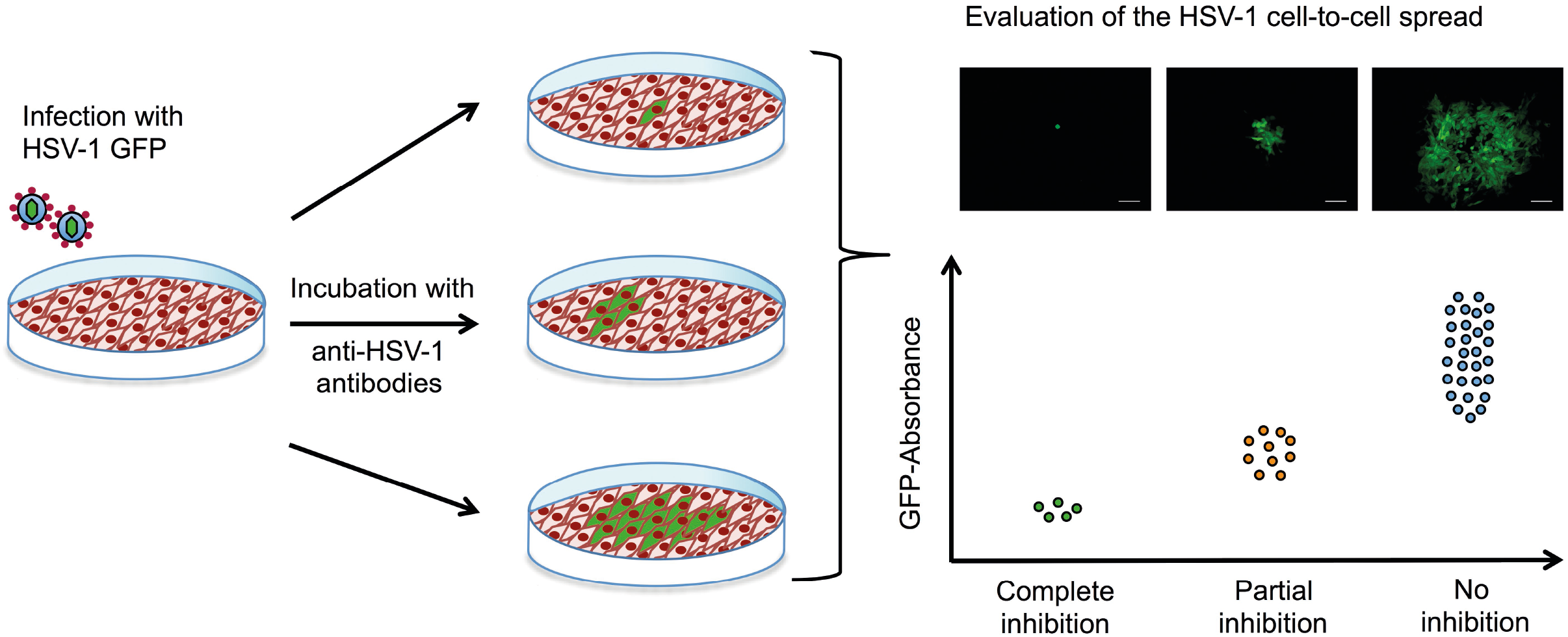
Assessment of the HSV-1 cell-to-cell spread inhibiting antibodies of human plasma or serum samples using the HSV-1 ΔgE GFP reporter virus-based screening method. The procedure was based on assessing the extent of plaque formation, which was proportional to the GFP-signal emitted by the infected cells. Confluent Vero cells were infected with HSV-1 ΔgE GFP reporter virus at low MOI. Infected cell cultures were overlaid with a medium containing either sera or plasma samples from HSV-seropositive humans at a 1:40 dilution. After 72 h hours of incubation, plaque formation was qualitatively assessed by fluorescence microscopy and simultaneously the GFP-signal was quantified as relative fluorescence units (RFU).

### Evaluation of an HSV-1-ΔgE-GFP-based high-throughput screening assay for cell-to-cell spread inhibiting antibodies

The HSV-1-ΔgE-GFP-based high-throughput screening assay was first evaluated using the humanized antibody mAb hu2c that completely inhibits HSV-1 and HSV-2 cell-to-cell spread (Fig. 2). Confluent Vero cell cultures were infected with HSV-1-ΔgE-GFP and subsequently overlaid with medium containing graded concentrations (0 - 500 nM) of mAb hu2c. Plasma or serum from an HSV-1 and HSV-2 double seronegative donor was added at a 1:40 dilution. Complete inhibition of the cell-to-cell spread could be observed at mAb hu2c concentrations between 125 and 500 nM (Fig. 2A). Almost complete inhibition was observed at 62.5 nM. At this concentration, only very small plaques with a maximum of 4 infected cells/plaque were visible (Fig. 2A) and the quantitative analysis showed an almost unchanged GFP signal compared to higher mAb hu2c concentrations (Fig. 2B). This concentration represents the lowest mAb hu2c concentration that almost completely inhibits the cell-to-cell spread (Fig. 2A, dashed line). At concentrations between 2 and 7.8 nM of mAb hu2c there was no visible reduction of the cell-to-cell spread (Fig. 2A) and the GFP-signal was notably higher when compared to concentrations above 62.5 nM mAb hu2c (Fig. 2B). Interestingly, plaques were smaller at mAb hu2c concentrations between 15.6 and 31.3 nM mAb hu2c (Fig. 2A) accompanied by only slightly increased GFP-signals (Fig. 2B), indicating a partial inhibition of the cell-to-cell spread at these concentrations.

**Figure 2:**
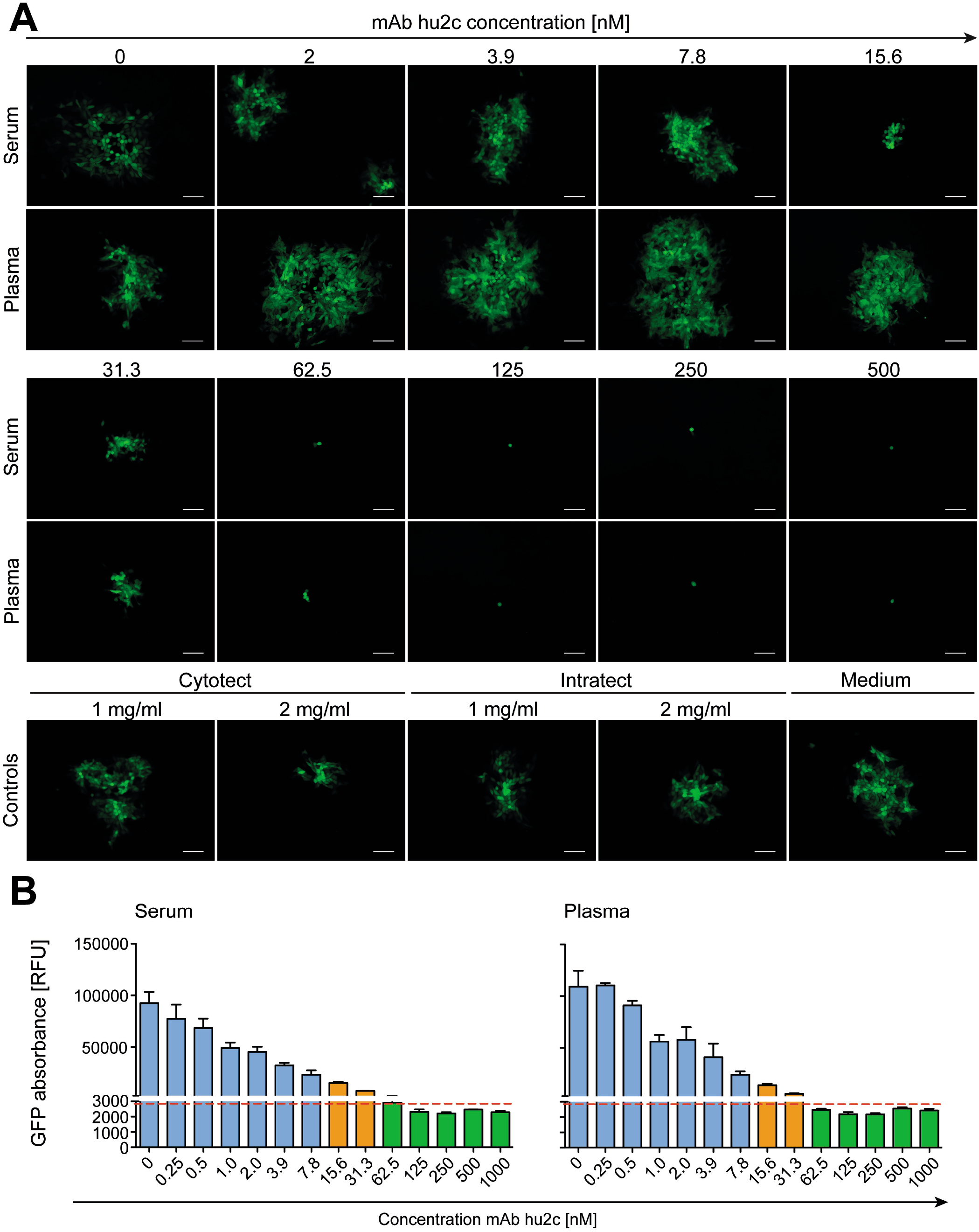
Evaluation of the HSV-1 cell-to-cell spread inhibition by mAb hu2c. The performance of the HSV-1 ΔgE GFP reporter virus-based assay was evaluated for the screening of sera and plasma that contain various concentrations of HSV-1 cell-to-cell spread inhibiting antibodies. (A) Confluent Vero cells growing on 24-well plates were infected with 200 TCID_50_/500 μl of the HSV-1 ΔgE GFP reporter virus. After 2 h of incubation, the inoculating medium was removed and the cells were overlaid with a medium containing sera or plasma from a HSV-1 seronegative donor at a 1:40 dilution. Additionally, the monoclonal, HSV-1/2 cell-to-cell spread inhibiting antibody mAb hu2c was added at a final concentration ranging from 0 to 1000 nM. After 72 h hours, plaque formation, which indicates HSV-1 spread via the cell-to-cell spread, was qualitatively assessed by fluorescence microscopy. 100x magnification, scale bar = 100 μm. (B) Additionally, the cell cultures were transferred to 96-well plates to quantify the GFP-signal as relative fluorescence units (RFU). Dashed line = cell-to-cell spread inhibiting concentration of mAb hu2c. Green bars = complete inhibition, orange = partial inhibition, blue = no inhibition of HSV-1 cell-to-cell spread.

These data clearly show that the quantitative measurement of the GFP signal correlates with the plaque expansion observed in cell-culture, which obviously represents the extent of the HSV-1 cell-to-cell spread. Furthermore, our assay distinguishes complete, partial and no inhibition of the HSV-1 cell-to-cell spread.

### Identification of HSV-1 elite responders with cell-to-cell spread inhibiting antibodies

To investigate whether humans can produce HSV-1 cell-to-cell spread inhibiting antibodies, plasma samples from 2496 blood donors were screened for cell-to-cell spread inhibiting properties. All samples were tested using the high-throughput screening assay described above. A mAb hu2c positive control was included on each 24-well plate. The efficacy of the plasmas regarding cell-to-cell spread inhibition was determined by dividing the GFP-signal of a culture containing the plasma of interest through the GFP-signal of mAb hu2c-treated control exhibiting “complete inhibition”. This quotient was termed inhibitory quotient (IQ) and represents the x-fold value of the GFP-signal measured for mAb hu2c. Plasmas were stratified according to their cell-to-cell spread inhibiting activity as completely inhibiting (IQ ≤ 1.5), partially inhibiting (IQ = 1.51 to < 2.8) and non-inhibiting (IQ ≥ 2.8). In total, 128 (5.1 %) of the plasmas showed complete and 1061 (42.5 %) exhibited partial inhibition (Fig. 3). The fold change of the remaining 1307 (52.4 %) plasmas was in a similar range as plasmas derived from HSV-1 seronegative donors and had no effect on the cell-to-cell spread. None of the 147 HSV-1 seronegative control plasmas exhibited partial or complete cell-to-cell spread inhibition, demonstrating the specificity of the assay.

**Figure 3:**
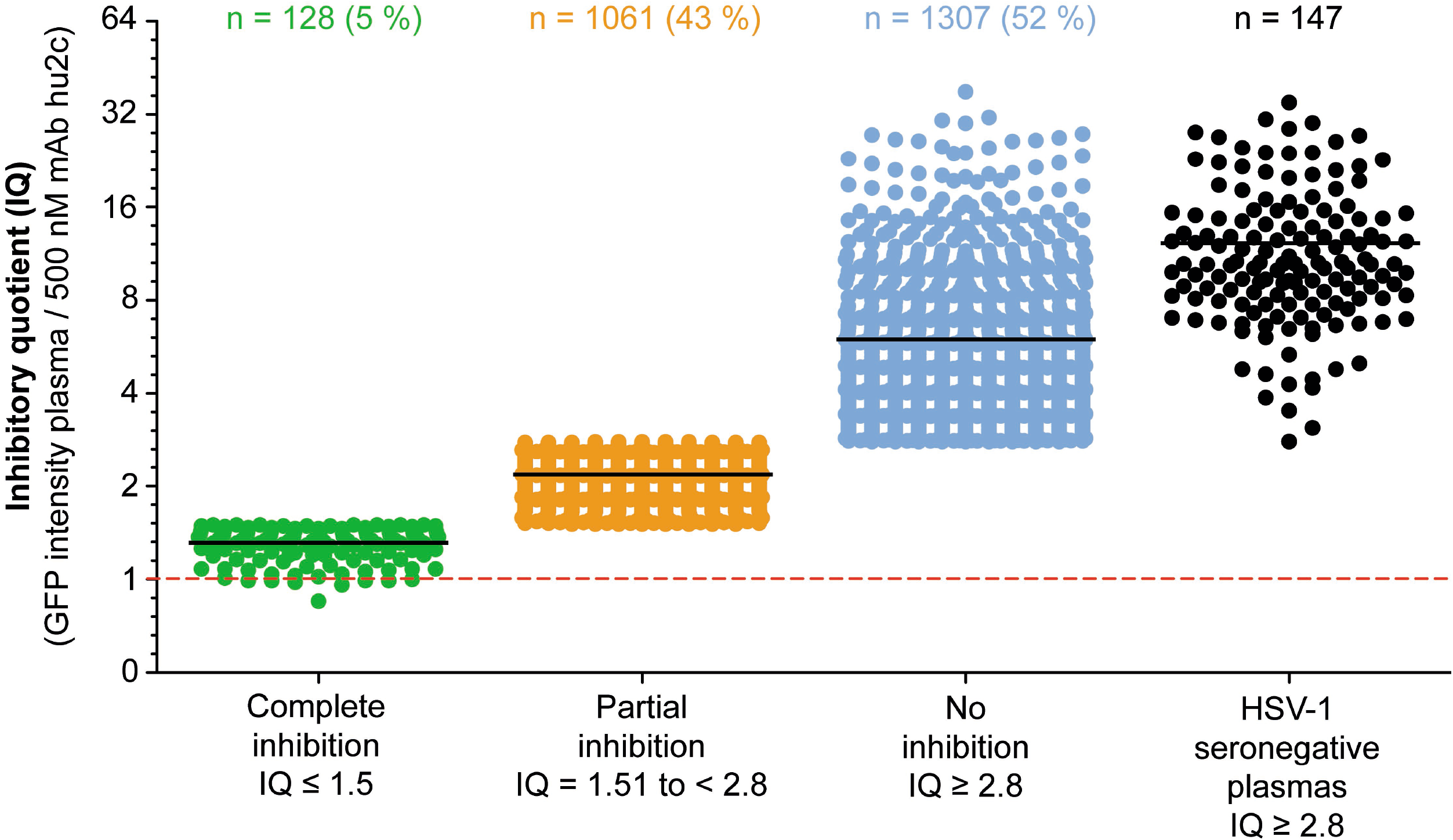
Assessment of the HSV-1 cell-to-cell spread inhibition capacity of plasma samples from HSV-1 seropositive blood donors. A total number of 2496 plasma samples from blood donors were investigated for HSV-1 cell-to-cell spreading properties using a HSV-1 ΔgE GFP reporter virus-based assay as described above. The efficacy of the plasma samples regarding cell-to-cell spread inhibition is shown as a fold change of the 500 nM mAb hu2c threshold (dashed line). At this concentration mAb hu2c completely inhibits the HSV-1 cell-to-cell spread. The efficacy of the plasma samples regarding cell-to-cell spread inhibition was determined by dividing the GFP-signal of a cell culture treated with a plasma sample through the GFP-signal of mAb hu2c control. This quotient was termed inhibitory quotient (IQ) and represents the x-fold value of the GFP-signal measured for mAb hu2c. The plasma samples were classified as completely cell-to-cell spread inhibiting (green dots, IQ ≤ 1.5), partially inhibiting (orange dots, IQ = 1.51 - 2.8) and non-inhibiting (blue dots, IQ ≥ 2.8). Each point represent the IQ for each donor, horizontal bars represent the median value.

### Assessment of the frequency of HSV reactivations in plasma donors

Next, we assessed the frequency of HSV reactivations in blood donors to investigate whether there is a correlation between the presence of cell-to-cell spread inhibiting antibodies and the frequency of reactivations. For this purpose, we conducted a retrospective survey including 158 blood donors that were randomly selected from the complete inhibition group (n = 47), partial inhibition group (n = 58) and the no inhibition group (n = 53). The HSV-seropositive status was confirmed by a diagnostic IgG ELISA. The biometric characteristics of the three different inhibition groups are summarized in Table 1. All three groups were comparable regarding mean age, gender, smoking behavior as well as the mean body mass index (BMI). Next, the three different groups were evaluated regarding the frequency of HSV reactivations. The frequency of HSV reactivations was recorded according to the observed occurrence of oral or genital lesions with less than one time per year or one or more symptomatic reactivations per year (< 1 or ≥ 1 reactivation per year). Interestingly, study participants from the complete inhibition group showed significant lower frequencies of HSV reactivation as compared to the groups that show only partial or no cell-to-cell spread inhibition capacity (Fig. 4). Only 17 % of individuals from the complete inhibition group reported one or more reactivation per year, whereas the frequency of at least one reactivation per year was 38 % in the partial inhibition group and 36 % in the no inhibition group. These data clearly demonstrate a significant correlation between the presence of cell-to-cell spread inhibiting antibodies and a lower rate of HSV-reactivation, providing a strong argument for their functional relevance in preventing recurrent herpes disease.

**Figure 4:**
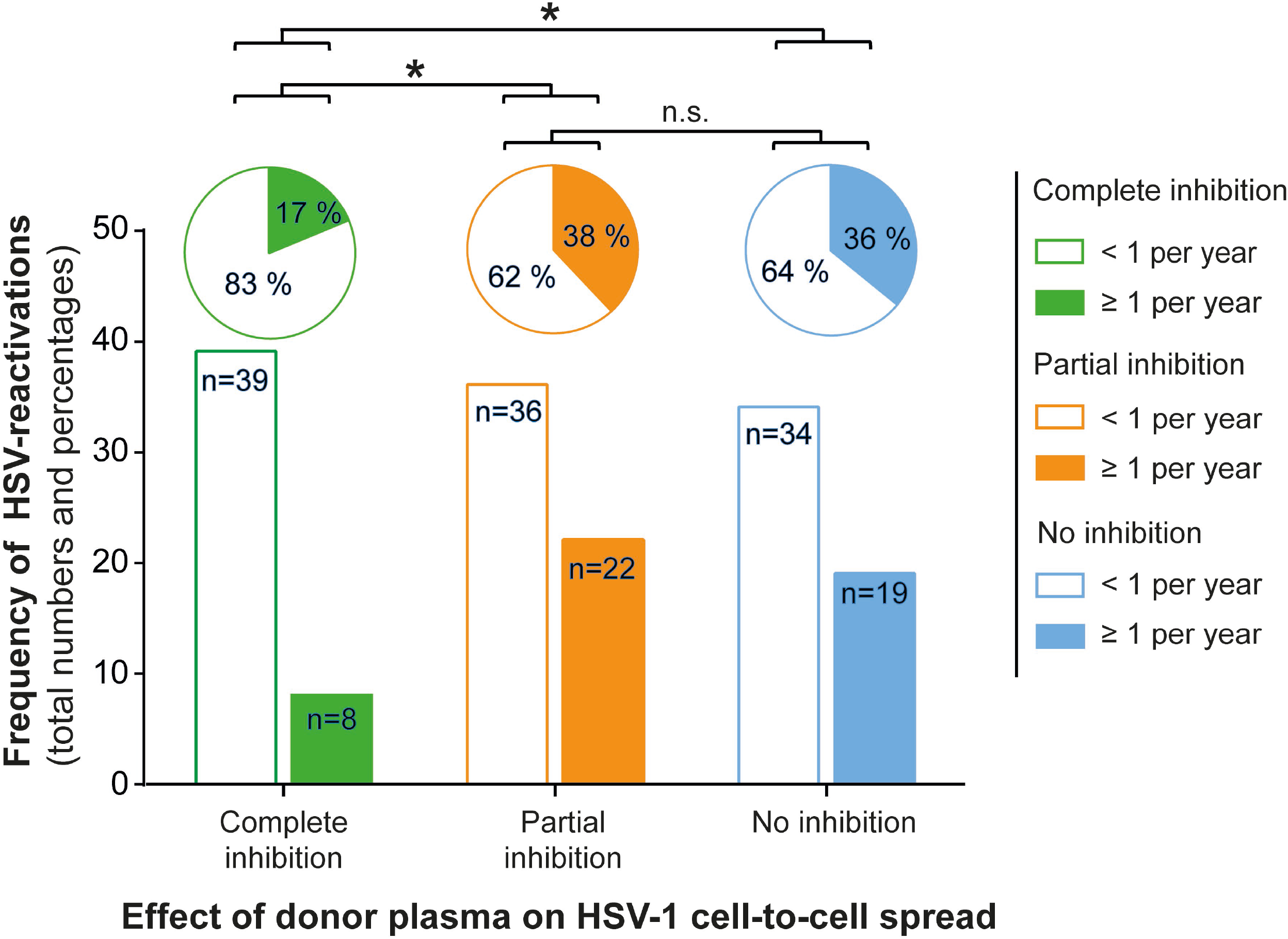
Correlation between protective antibody response and the frequency of HSV reactivation. A total number of 158 HSV seropositive blood donors previously being tested for cell-to-cell spread inhibiting antibodies were retrospectively interviewed for the frequency of symptomatic HSV reactivations per year. The donors were divided into the three groups (complete inhibition, n = 48; partial inhibition, n = 58 and no inhibition, n = 53) according to the performance of the donor plasmas on the HSV-1 cell-to-cell spread inhibition. The total numbers of donors are depicted as a bar chart and the percentages as a pie chart above. Differences in the annual frequency of HSV reactivation were analyzed using the Fischer’s exact test. Significant changes (*\*p* < 0.05) are indicated by asterisks and non-significant changes (*p* > 0.05) are labeled as “n.s.”.

### Determination of the neutralizing antibody titers of blood donor plasmas

To investigate whether antibodies capable to neutralize HSV-1 but incapable to block the cell-to-cell spread of the virus have an impact on the frequency of HSV-1 reactivations, we determined the neutralizing titers of the donor plasmas that completed the survey.

Although there was a significant difference in the frequency of reactivations between the complete inhibition group and partial inhibition group (Fig. 4), the neutralizing antibody titers were at similar levels (Fig. 5). Only the no inhibition group was shown to have significantly lower neutralization titers compared to the complete inhibition and partial inhibition group. These results indicate that the significantly lower HSV-reactivation likelihood observed for the complete inhibition group correlates with cell-to-cell spread inhibiting antibodies but not with antibodies that neutralize the virus but fail to counteract the cell-to-cell spread.

**Figure 5:**
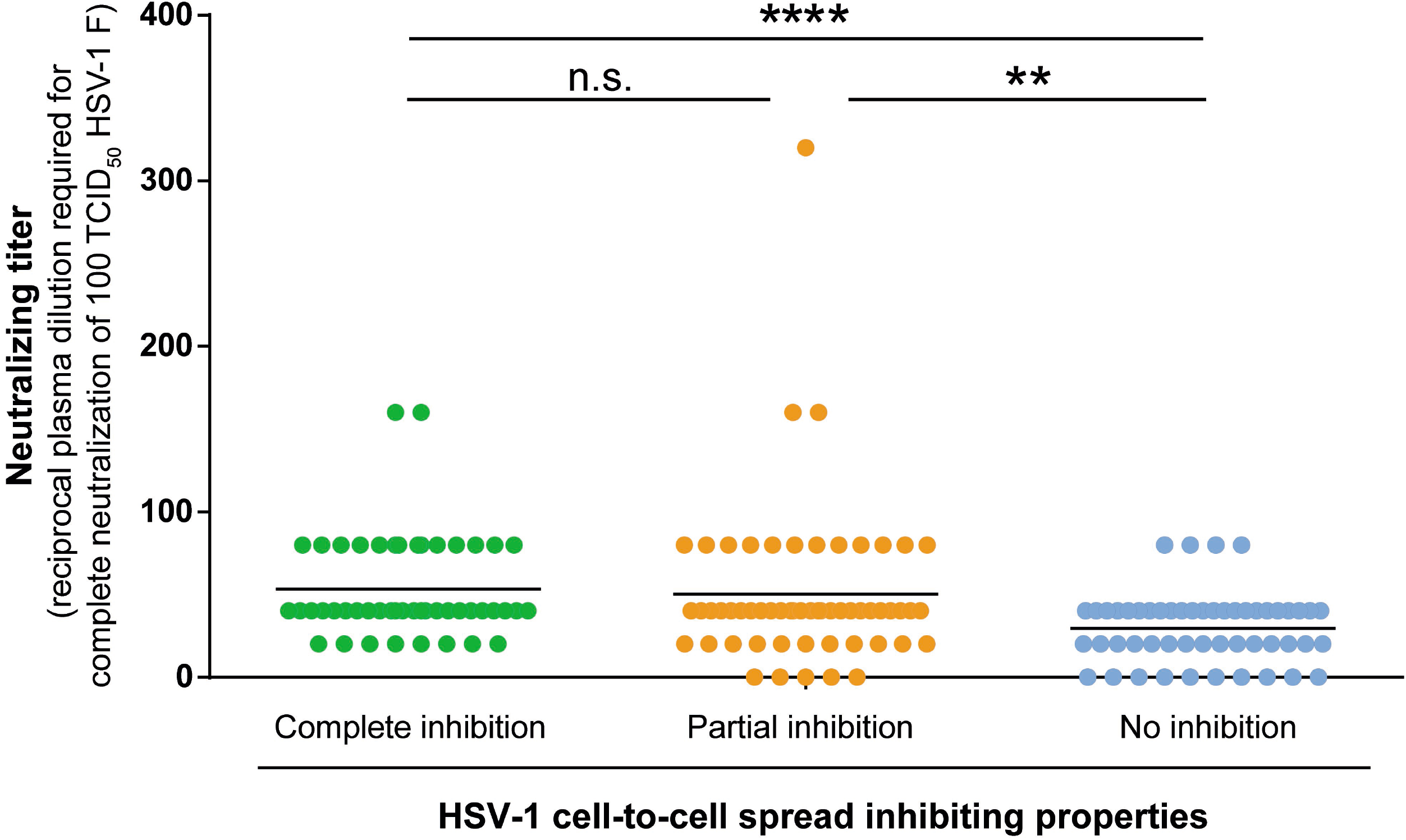
HSV-1 neutralizing antibody titers of the plasma samples of the three inhibition groups (complete, partial and no inhibition group). Serial dilutions of the respective plasma samples (1:20 to 1:2560) were preincubated with 100 TCID_50_ HSV-1 F for one hour and subsequently added to Vero cells in 96-well microtiter plates. After 48 h of incubation, the cytopathic effect was analyzed and the respective neutralization titers were determined. Data sets were statistically analyzed using the One-way ANOVA followed by the Dunn’s multiple comparison *post-hoc* test. Significant changes (\*\**p* < 0.01, \*\*\*\**p* < 0.0001) are indicated by asterisks and non-significant changes (*p* > 0.05) are labeled as “n.s.”.

In conclusion, we showed for the first time that (i) about five percent of HSV-1 seropositive blood donors (elite responder) are able to produce HSV-1 cell-to-cell spread inhibiting antibodies and (ii) that the presence of these antibodies correlate with a significantly lower risk of HSV-reactivation.

## Discussion

In the present study, we investigated whether humans are able to produce antibodies that effectively block the HSV-1 cell-to-cell spread upon natural infection. We demonstrated that humans are principally able to produce such cell-to-cell spread inhibiting antibodies against HSV-1.

In our large cohort of 2496 blood donors, we identified a small proportion of 128 (5.1 %) that had a sufficiently high antibody-concentration to block the cell-to-cell spread of HSV-1 in cell culture (elite responder). Most importantly, these individuals reported a significantly lower frequency of symptomatic HSV-reactivations when compared to people with lower or no detectable cell-to-cell spread inhibiting antibodies.

Interestingly, 42.5 % of the plasmas showed a partial inhibition of the cell-to-cell spread, indicating that there might be cell-to-cell spread inhibiting antibodies in the plasmas, but at lower concentrations. The average concentration of antibodies in human sera/plasma was described with 11 mg/ml [21]. In the present study, we tested the plasma samples at a 1:40 dilution, which corresponds to a median IgG concentration of 0.28 mg/ml. At least in 5 % of individuals whose plasmas showed complete cell-to-cell spread inhibition in our high-throughput, this concentration was sufficient to prevent reinfections.

However, we found that there was no correlation between neutralizing antibody titers and the frequency of HSV reactivation. Despite similar neutralizing antibody titers, people who had cell-to-cell spread inhibiting antibody concentrations in plasma reported significantly fewer rate of HSV-reactivations than people with insufficient concentrations of such antibodies did. These data provide evidence for the unique protective role of cell-to-cell spread inhibiting antibodies in HSV infection.

These data support prior findings. Neutralizing antibodies, which are not necessarily inhibiting the cell-to-cell spread, contributed to protect from a severe course of disease [22]. The presence of HSV-specific antibodies in HSV-infected mothers has been suggested to decrease the risk of acquisition of HSV-2 by newborns [23, 24]. Similar findings were reported in mice. Maternal antibodies were shown to access neural tissues of the fetus or neonate, thereby protecting neonatal mice against HSV-1 neurological infection and death [24]. Notably, in animal studies neutralizing antibodies blocking virus entry and cell-to-cell spread were superior to normal neutralizing antibodies that did not inhibit the cell-associated viral spread [20]. These data are in line with our here presented findings.

In conclusion, by using a high-throughput HSV-1-ΔgE-GFP reporter assay we have demonstrated that HSV-1 infected humans are able to produce cell-to-cell spread inhibiting antibodies. In addition, we were able to show that the presence of these antibodies directly correlates with a significantly lower frequency of HSV reactivations, representing a correlate of protection. Plasmas of these individuals may be used for passive immunization strategies. Isolation of the cell-to-cell spread inhibiting antibodies may contribute to develop novel antibody-based interventions for prophylactic and therapeutic use. Moreover, characterizing of epitopes recognized by these antibodies may contribute to optimize the target antigens for novel vaccine approaches.

## Material and methods

### Ethics statement

The study was performed in accordance with The Code of Ethics of the World Medical Association (Declaration of Helsinki) and was approved by the ethical committee of the University of Ulm and the University Hospital Essen.

### Antibodies, Sera and Plasma

Sera and plasma samples were harvested during routine blood donations at the Institute of Transfusion Medicine, University of Ulm, Germany. The humanized monoclonal antibody mAb hu2c was produced and purified as described previously [15].

### Viruses

HSV-1 strain F, HSV-2 strain G and HSV-1-ΔgE-GFP reporter virus were propagated in Vero cells and stored at −80 °C. HSV-1-ΔgE-GFP was kindly provided by Hartmut Hengel (Institute of Virology, Freiburg, Germany) and initially described by Farnsworth et al. [21]. Viral titers were determined by a standard endpoint dilution assay and calculated as 50 % tissue culture infectious dose (TCID_50_)/ml as previously described [22].

### Cells

Vero cells (American Type Culture Collection, ATCC, CCL81, Rockville, MD) were cultured in Dulbecco’s Modified Eagle Medium (DMEM, Life Technologies Gibco, Darmstadt, Germany) containing 10 % (v/v) fetal calf serum (FCS; Life Technologies Gibco), 100 U/ml penicillin and 0.1 mg/ml streptomycin.

### HSV-1-ΔgE-GFP based screening for cell-to-cell spread inhibiting antibodies

To investigate whether humans are able to produce cell-to-cell spread inhibiting antibodies, we established an HSV-1-ΔgE-GFP reporter virus based assay for the high-throughput screening of HSV-1 seropositive human serum or plasma samples. The assay was evaluated using the HSV-1 and HSV-2 cell-to-cell spread inhibiting antibody mAb hu2c [15].

Highly permissive Vero cells were seeded on 24-well plates at a density of 1 x 105 cells/well. Confluent cell cultures were infected with 200 TCID_50_ HSV-1-ΔgE-GFP/well (MOI = 0.001). After 2 hours of incubation, the inoculation medium was removed and the cell cultures were incubated with serial dilutions of mAb hu2c (0 - 500 nM). Commercial polyclonal human antibody preparations, Cytotect and Intratect (Biotest, Dreieich, Germany), were used as controls at a concentration of 1 or 2 mg/ml. To standardize the background levels, all purified antibodies were applied in medium containing serum or plasma from an HSV-1 and HSV-2 seronegative donor at a 1:40 (v/v) dilution. After 3 days of incubation, the plaque formation was examined by fluorescence microscopy (Axio Observer D1, Zeiss). Additionally, the fluorescence levels were quantified. For this purpose, the cell culture medium was removed, the cells washed with PBS, detached with Trypsin/0.5% EDTA (Life Technologies Gibco), resuspended and transferred to 96-well plates. GFP-signals were quantified using the Mithras^2^ LB 943 microplate multimode reader (Berthold Technologies).

### High throughput screening of plasmas for the inhibition of HSV-1 cell-to-cell spread

A total number of 2496 human plasmas were screened for the inhibition of the HSV-1 cell-to-cell spread using the high throughput assay as described above. Human plasma samples were applied at 1:40 dilutions. The monoclonal humanized antibody mAb hu2c served as positive control at a concentration of 500 nM (75 μg/ml) diluted in plasma from an HSV-1/2 seronegative donor (1:40 in cell culture medium). At this concentration of mAb hu2c, the HSV 1 cell-to-cell spread is completely blocked. After 72 hours of incubation, the GFP-signal was measured using the Mithras^2^ LB 943 microplate multimode reader (Berthold Technologies). Fluorescence-values for individual plasma samples were compared with the GFP-intensity measured for mAb hu2c for each plate. The values obtained for the plasma samples were then normalized to the mAb hu2c control and calculated as the x-fold value of the mAb hu2c signal.

### Identification of HSV-seropositive plasmas by ELISA

The HSV-seropositivity status of donors completing the survey was confirmed by ELISA using the anti-herpes simplex virus type 1 and 2 IgG human ELISA kit (abcam, Cambridge, United Kingdom) according to manufacturer’s instructions.

### Retrospective survey to determine the frequency of HSV-reactivations

To investigate the role of HSV-1 cell-to-cell spread inhibiting antibodies in HSV-seropositive people, we assessed the frequency of symptomatic HSV reactivations in the frame of a retrospective survey. The data acquisition comprised the annual numbers of symptomatic oral or genital HSV-reactivations characterized by the occurrence of characteristic lesions. Furthermore, data on general topics like age, gender, body mass index (BMI) as well as smoking behavior were collected. The survey enrolled a total number of 158 blood donors stratified in three comparable groups according to the presence of cell-to-cell spread inhibiting antibodies as complete inhibition (n = 47), partial inhibition (n = 58) and no inhibition (n = 53).

### Neutralization assay

To investigate the neutralizing antibody titers in blood donor plasmas, a neutralization assay was performed as previously described [16]. Briefly, serial dilutions (1:20 to 1:2560) of the respective plasma samples were pre-incubated with 100 TCID50 of HSV-1 F for one hour at 37 °C and added afterwards to confluent Vero cells cultured in 96-well microtiter plates. After 72 hours, the cytopathic effect was analyzed by microscopy and the reciprocal neutralization titer was determined.

## Acknowledgements

The authors thank Delia Cosgrove for the proofreading of the manuscript.

## Author Contributions

SW, MA, RD, MD and UWA performed the experiments. SW, MA, RD, MD, UWA, BG, MR, UD, OW, MT, CH and AK analyzed the data. KR conducted the survey. RL, MR and AK planned the study. SW, MA, CH and AK wrote the manuscript. All authors approved the final version of the manuscript.

